# Stable association of a Drosophila-derived microbiota with its animal partner and the nutritional environment upon transfer between populations and generations

**DOI:** 10.1101/111492

**Authors:** Mélisandre A. Téfit, Benjamin Gillet, Pauline Joncour, Sandrine Hughes, François Leulier

## Abstract

In the past years, the fruit fly *Drosophila melanogaster* has been extensively used to study the relationship between animals and their associated microbes. Compared to the one of wild populations, the microbiota of laboratory-reared flies is less diverse, and comprises fewer bacterial taxa; nevertheless, the main commensal bacteria found in fly microbiota always belong to the *Acetobacteraceae* and *Lactobacillaceae* families. The bacterial communities associated with the fly are environmentally acquired, and the partners engage in a perpetual re-association process. Adult flies constantly ingest and excrete microbes from and onto their feeding substrate, which are then transmitted to the next generation developing within this shared habitat. We wanted to analyze the potential changes in the bacterial community during its reciprocal transfer between the two compartments of the niche (i.e. the fly and the diet). To address this question, we used a diverse, wild-derived microbial community and analyzed its relationship with the fly population and the nutritive substrate in a given habitat. Here we show that the community was overall well maintained upon transmission to a new niche, to a new fly population and to their progeny, illustrating the stable association of a Drosophila-derived microbiota with its fly partner and the nutritional environment. These results highlight the preponderant role of the nutritional substrate in the dynamics of Drosophila/microbiota interactions, and the need to fully integrate this variable when performing such studies.

## 1. Introduction

Thanks to its ease of manipulation and genetic tractability, the fruit fly *Drosophila melanogaster* has been used as a model organism for more than a century (Kohler 1994; Sang 2001). Like all other animal species, Drosophila have been living and evolving in close association with microorganisms (McFall-Ngai et al. 2013), and such partnership impact various traits of the fly partner’s physiology including growth, developmental timing, stress resistance, immune response, metabolism, lifespan and behavior (Brummel et al. 2004; Ryu et al. 2008; Sharon et al. 2010; Shin et al. 2011; Guo et al. 2014; Petkau et al. 2014; Venu et al. 2014; Wong et al. 2014; R. I. Clark et al. 2015; Téfit & Leulier 2017). Most of the functional studies on Drosophila-microbiota interaction are based on manipulating gnotobiotic animals generated through the association of germ-free animals with one to five cultured commensal bacterial strains. Although not as complex as the one of mammals, the microbiota of laboratory-reared Drosophila generally comprises up to twenty community members (Broderick & Lemaitre 2012; Erkosar et al. 2013). The exact microbiota composition may vary across studies, but some common features dominate. For example, the represented species differ, but the community diversity is quite low at the higher taxonomic levels and the most represented bacteria always belong to the *Acetobacteraceae* and *Lactobacillaceae* families (Staubach et al. 2013; Ma et al. 2015). Furthermore, analyses of the communities associated with wild-caught Drosophila populations confirmed the low diversity of bacterial taxa identified. Indeed, *Enterobacteriaceae* and *Acetobacteraceae* families, as well as the *Lactobacillales* order represent the major components of the "wild" flies microbiota (Chandler et al. 2011; Staubach et al. 2013). In addition, many rare taxa are found in wild populations (such as *Erwinia, Pantoea* or *Gluconobacter)* and their identity vary among studies (Chandler et al. 2011; Staubach et al. 2013).

In the wild, *Drosophila melanogaster* lives and feeds on rotting fruits, which represent an eminently microbe-rich environment. Fruit flies thus constantly ingest and excrete microorganisms which in turn (re-)colonize the niche and will then be transmitted to the next generation (Erkosar et al. 2013). In laboratory settings, the situation is similar since flies are reared in vials, a closed environment in which this colonization cycle also takes place. With the advance of the Drosophila microbiota research field, the idea of a resident, stable and defined microbiota of the fly has been challenged (Wong et al. 2013). Indeed, so far there is no published evidence supporting the existence of bacterial species that persistently reside within the fly gut, and are different than the ones encountered in the immediate environment of the animal. The relationship between the fly and its microbiota appears to be more transient and highly dependent on the nutritive substrate on which Drosophila develops and lives (Sharon et al. 2010; Chandler et al. 2011; Staubach et al. 2013). These observations highlight the importance to consider the fly niche as a whole (i.e. including its nutritional substrate) when studying the interaction between the fruit fly and its associated bacteria.

To investigate the relationship between Drosophila, its microbiota and the nutritional substrate, we surveyed the dynamics of the structure and composition of a bacterial commensal community. We wanted to analyze the potential changes in the bacterial community during its reciprocal transfer between the two compartments of the niche (i.e. the fly and the diet). Ultimately, we were interested in understanding whether the environmental niche comprises one common bacterial community shared between both compartments, or rather sub-communities associated with either the flies or the nutritive substrate. To this end, we established the profile of the bacterial communities associated with flies and with their diet and observed that the flies did not seem to actively select for or against specific bacterial orders or families. Indeed, despite minor fluctuations in the bacterial taxa representation, there was a high degree of similarity between the composition of the bacterial community associated with the flies and the one of the community in the diet. Additionally, the community was overall well maintained upon transmission to a new habitat, to a new fly population and to their progeny. Taken together, the results of this study illustrate the stable association of a Drosophila-derived microbiota with both its animal partner and the nutritional environment and highlight the need to take into account the role of the diet when studying the interaction between Drosophila and its microbiota.

## 2. Material and methods

### 2.1. Fly stocks and husbandry

Laboratory-reared and wild-caught Drosophila populations were used in this study, both carrying the bacterial endosymbiont Wolbachia. Laboratory-reared *y,w* flies were kept on a standard yeast/cornmeal diet containing for 1L: 50g inactivated yeast (Bio Springer, Springaline BA95/0-PW), 80g cornmeal (Westhove, Farigel maize H1), 10g agar (VWR, ref. #20768.361), 5.2g methylparaben sodium salt (referred to as Moldex, MERCK, ref. #106756) and 4ml 99% propionic acid (CARLO ERBA, ref. #409553). Wild-caught flies were collected from rotten tomatoes in a garden in Solaize (France) and reared on a yeast-sucrose diet devoid of chemicals (YS-), and containing for 1L: 15g inactivated yeast, 25g sucrose (Sigma Aldrich, ref. #84100), 80g cornmeal and 10g agar. For the experiments, this diet was supplemented with 2.5ml 99% propionic acid and the quantity of yeast was decreased to 10g/L (YSexp). All experimental flies were kept in incubators at 25°C, with a 12h/12h light/dark cycle.

### 2.2. Generation of axenic Drosophila stocks

To generate axenic flies, eggs were collected overnight and treated in sterile conditions with successive 2 minutes baths of bleach and 70% ethanol. Bleached embryos were then rinsed in sterile water for another 2 minutes and placed on sterile standard diet supplemented with an antibiotic cocktail (50μg ampicillin, 50μg kanamycin, 50μg tetracycline and 15μg erythromycin per liter of fly diet). Emerging adults were tested for axenicity by crushing and plating of the fly lysate on different bacterial culture media. The absence of Wolbachia contamination in the axenic stocks was confirmed by PCR, using the following general or strain specific primer pairs:

- WolbFWD: 5’ - TGGTCCAATAAGTGATGAAGAAAC - 3’
- WolbREV: 5’ - AAAAATTAAACGCTACTCCA - 3’
- WSP81FWD: 5’ - TTGTAGCCTGCTATGGTATAACT - 3’
- WSP691REV: 5’ - GAATAGGTATGATTTTCATGT - 3’

Germ-free flies were kept on antibiotic diet for a few generations and conventionally reared stocks were used to generate new axenic stocks regularly.

### 2.3. Wild microbiota inoculation and samples collection

Twenty-five males from the wild-caught population were put in a cage to seed sterile YSexp diet (contained in a Ø60mm petri dish) with their microbiota. After 4 days, fly and diet samples were retrieved from the cage and treated separately: on one hand, 4 replicate groups of 10 flies were crushed in 500μL sterile PBS and on the other hand, for each cage, 3 replicates of 250mg of microbes-seeded diet were crushed in 1mL of sterile PBS. 50μL of each diet resuspension replicate was collected in order to assess the microbial diversity of the diet at the beginning of the experiment. These aliquots were pooled and 50μL of the resulting mix were then used to inoculate fresh sterile YSexp diet in a Ø1.5cm fly tube. 5 to 10 days old axenic *y,w* adults were added in these tubes and left to lay eggs on the inoculated diet. After 4 days, the ex-axenic adults were collected and crushed in sterile PBS. Their progeny was then left to develop on the wild microbiota-inoculated YSexp diet and after two weeks triplicate fly and diet samples were collected. The whole experiment was conducted in duplicate (**Figure 3**).

### 2.4. DNA extraction, amplification and sequencing

Using the UltraClean^®^ Microbial DNA Isolation Kit (MO BIO Laboratories, Carlsbad, CA, USA), DNA was isolated from all the samples collected during the wild microbiota inoculation experiment following the manufacturer’s instructions. For experiments on the conventionally reared *y,w* stock, DNA was isolated from 5 to 10 days old flies with a protocol adapted from (Wong et al. 2013). Groups of 10 flies (5 females + 5 males) were homogenized in 300μL lysis buffer (20mM Tris-HCl, pH8, 2mM sodium EDTA, 1.2% Triton-X100, 20mg/mL lysozyme) by bead beating on a Precellys24 Sample Homogenizer (Bertin Instruments; 6500rpm, 2x30 seconds) and incubated at 37°C for 90 minutes, with another round of bead-beating at 45 minutes. 300μL 2X extraction buffer (400mM Tris-HCl, pH8.5, 500mM NaCl, 50mM EDTA) were added, together with 20μL 20% SDS and 15μL proteinase K (20mg/mL). Samples were then incubated overnight at 42°C, fly tissues debris were removed by phenol-chloroform treatment and DNA precipitated with 1:10 volume of 3M sodium acetate. The supernatant was mixed with 2.5 volumes of ice-cold 100% ethanol and incubated at −20°C for 15 minutes before centrifugation at 4°C for 30 minutes and at 15000g. After discarding the supernatant, each pellet was washed in 1mL ice-cold 70% ethanol, dried and resuspended in 20μL low TE buffer. The variable region V3 of the 16S rRNA bacterial gene was amplified by PCR using the primers 338F and 700R (Wang & Qian 2009). Barcode sequences, taken from Hamady et al. (2008), were added 5’ of each reverse and forward primers for subsequent multiplexing of the samples in the sequencing approach **(Table 1)**. PCR products were processed for sequencing with the Ion Torrent™ Personal Genome Machine^®^ (PGM) system (Thermo Fisher Scientific Inc.). PCR mixes were prepared in a dedicated room where no DNA is manipulated and DNA extracts were subsequently added in a room where no amplified DNA is present. Both rooms were decontaminated by UV lights and all surfaces under hoods were cleaned with DNA-ExitusPlus (PanReac, AplliChem) between each work session. Various controls were added to the experiments to monitor possible bacterial contamination at different steps (mock extraction, negative and aerosol PCR controls). PCR reactions were performed in 25 *μ*l using the Environmental Master Mix 2.0 (Thermo Fisher Scientific) and 0.25*μ*M of each primer with the following PCR program: 10 minutes at 94°C, 35 cycles at 94°C 40 seconds, 55°C 40 seconds and 72°C 1 minute, with a final extension step à 72°C for 7 minutes. PCR product purification was carried out according to the manufacturer (Nucleospin Gel and PCR Clean-up, Macherey Nagel) and amplicons were eluted in 30*μ*l of NE buffer. Equimolar amounts of the purified amplicons were used to create a unique library using the protocol Preparing Short Amplicon (<350pb) Libraries Using the Ion Plus Fragment Library Kit. Quantitation and quality assessment of library was performed on 2200 Tapestation analyzer using the High Sensitivity D1000 ScreenTape kit (Agilent Technologies). The library was subsequently processed with the Ion PGM Template OT2 HiQ 400 Kit and sequenced with the Ion Torrent PGM on a 316v2 chip using the Ion PGM HiQ Sequencing Kit.

**Table 1.**
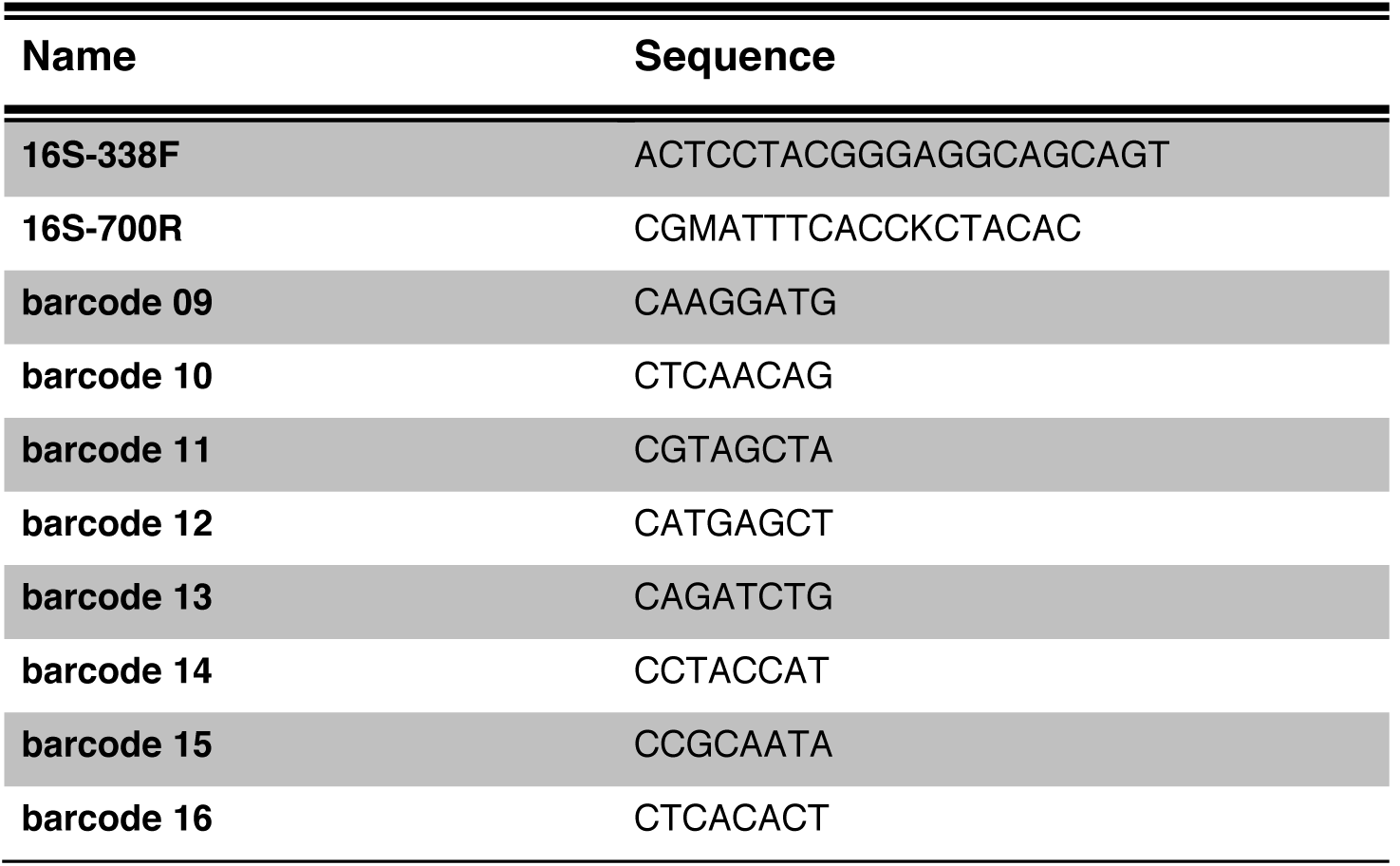
Sequences of primers and barcodes used for the PCR amplification and sample identification in 16S-based community profiling protocols.

### 2.5. Processing of the sequencing data and analysis

Reads obtained from the sequencing were demultiplexed and sorted by sample of origin based on the barcodes used. Using the Cutadapt software (Martin 2011), reads were then trimmed to remove barcodes and primers sequences and those inferior to 200bp were discarded. At this point, all reads were merged into a single fasta file for downstream analyses, with their sample identification available in a separate group file. Pooled reads were processed with the Mothur program, version 1.34.4 (Schloss et al. 2009) following the pipeline *Ion Torrent sequence analysis using Mothur* contributed to the Mothur community by Sukithar Rajan. Reads were aligned against the SILVA database alignment file (release 123). Alignment result was screened and poorly aligned sequences were filtered out. Chimeric sequences were also identified and removed from the file before classification with the SILVA taxonomic outline. Taxonomy files were then analyzed using custom R scripts to calculate the frequencies of each taxonomic level of interest in the samples. Additionally, when present, the frequencies of taxons observed in the controls samples were subtracted from the corresponding frequencies in the experimental samples. The controls correspond to the sequencing of DNA extraction and PCR amplification reagents, without any tissue or DNA added. They were processed together with the experimental samples to monitor the potential environmental contaminations often encountered in this type of studies (Salter et al. 2014). The frequency tables were then used to generate barplot graphs describing the taxonomic composition of the samples. Only taxa for which the frequency is above 1% are represented on the graphs.

### 2.6. Statistical analyses

Data were analyzed using R and Rstudio (versions 3.1.2 and 0.98.1091 respectively), to generate barplot graphs. For the statistical analyses, type III ANOVAs have been performed to test the combined effect of the three factors on alpha-diversity and proportion of bacteria. When the effects were significant, pairwise-t-tests were used to compare between combinations of modalities. To cluster the samples based on similarities in bacteria composition, principal component analyses and hierarchical clustering have been performed with the FactoMineR package. The number of clusters for the hierarchical clustering has been chosen based on a drop in inertia gain.

## 3. Results

### 3.1. Low diversity of the bacterial community associated with a Drosophila stock

To determine whether the diversity of the community associated with our laboratory reference stock *yellow,white (y,w)* was suitable for a community dynamics study, we first assessed its composition. We performed the profiling of the microbiota of conventionally reared *y,w* flies with 16S-based sequencing. Briefly, the V3 variable region of the bacterial 16S ribosomal RNA gene was amplified and the resulting amplicons were sequenced with an Ion Torrent™ PGM system. We obtained over 2.5 millions of reads that were then processed to identify the bacterial taxa (see section 2. Material and Methods section for a detailed description of the sequencing protocol and analysis pipeline). The sequencing results showed a preponderance of the *Lactobacillales* order, together with a minor representation of *Rickettsiales* and *Corynebacteriales* **(Figure 1A)**. This low diversity of bacterial orders was further confirmed at the family and genus levels **(Figure 1B and C)**. Indeed, the *Lactobacillus, Wolbachia* and *Corynebacterium* genera were almost the sole representatives of their respective orders. Only in the case of *Lactobacillales* was there another member of the order represented; *Enterococcus* bacteria were also present in the samples but at a much lower frequency **(Figure 1C)**. This description is based on the analysis of three biological replicates pooled together. It nonetheless holds true when the three replicates are analyzed separately **(Figure S1)**.

**Figure 1.**
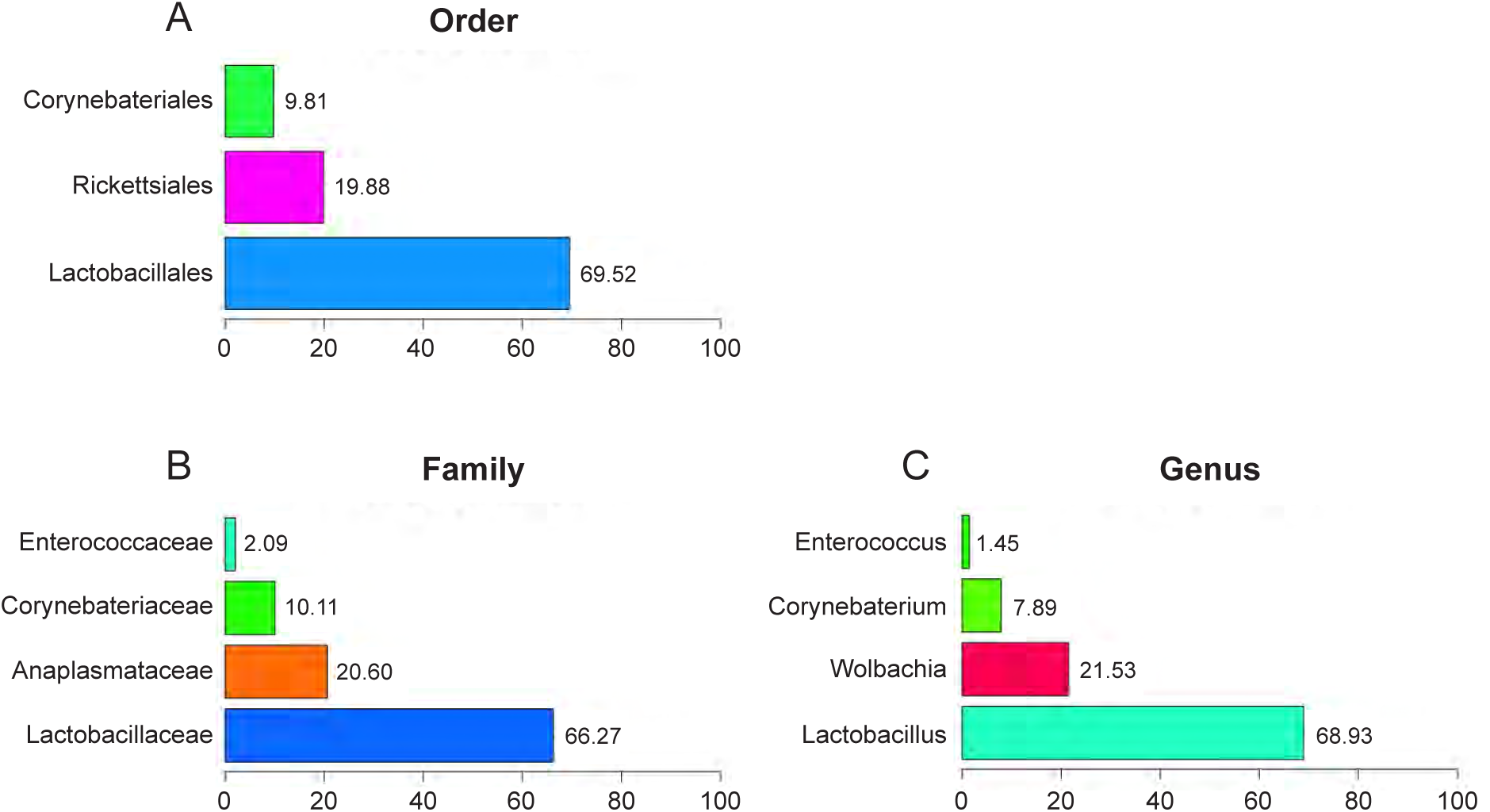
Community profile of the *yellow white* stock microbiota. Frequency of bacterial taxa present in the samples (pooled replicates). Taxonomic levels used: Order (A), Family (B) and Genus (C). Equivalent percentages are indicated next to the corresponding bars. Graphs only represent the taxa present at a frequency of 1% or above.

### 3.2. Preservatives contained in the fly diet dramatically impact the microbiota diversity

Given the very low complexity of the bacterial community associated with our conventionally reared *y,w* stock, we decided to search for a source of microbiota for our community dynamics study. Ideally, the starting bacterial community should be as diverse as possible to study the potential composition shifts occurring during its transfer between populations and generations. To maximize our chances to find flies with a diverse microbiota, we turned to the wild and collected Drosophila from rotting tomatoes. This population was composed of mostly *Drosophila melanogaster,* but the sorting of the flies was based just on physical features of *D. melanogaster* that are readily identifiable under a dissecting scope; therefore we cannot completely rule out the possibility that other visually resembling species were also present in the wild-derived population.

In the laboratory, we kept the wild-derived population on a preservative-free diet containing inactivated yeast and sucrose (YS- diet), a diet designed to favor the maintenance of bacterial diversity. In our initial effort to associate the wild-derived microbiota with axenic *y,w* eggs, we systematically faced invasive microbial contamination of the medium (likely of fungal origin) and subsequent death of the embryos and/or larvae. We therefore decided to reintroduce preservatives to the experimental diet, in a parsimonious manner. Our laboratory diets usually contain two chemicals serving as preserving agents: propionic acid and methylparaben sodium salt, referred to as Moldex (see section 2. Material and Methods section for detailed composition of the fly diet). To determine which of these two chemicals have the lowest impact on the bacterial diversity, we prepared YS- diets supplemented with propionic acid, Moldex or both. Wild-derived flies were placed on these diets and after several generations, a few adults from each sub-population were crushed and their lysate spread on different bacterial culture media. We observed that both chemicals prevented the invasive microbial contamination in the diet. However, the nature of the chemicals used in the diet fed to the flies had a major impact on the diversity of their microbiota **(Figure 2)**. Indeed, compared to that of flies reared on the YS- diet, the microbial diversity associated with flies fed a diet containing both propionic acid and Moldex was greatly reduced, as indicated by the low number and uniform shapes of the colonies growing on the plates. Between the two mono-chemical diets, the one containing only propionic acid seemed to preserve a better diversity than the Moldex-only one; the profile of microbial communities growing on plate after propionic acid-only treatment was more similar to the one obtained from the YS- diet. For subsequent experiments we therefore used a yeast/sucrose diet containing a low dose (2,5ml/L as compared to 4ml/L in our regular fly diet) of propionic acid as preservative (YSexp diet).

**Figure 2.**
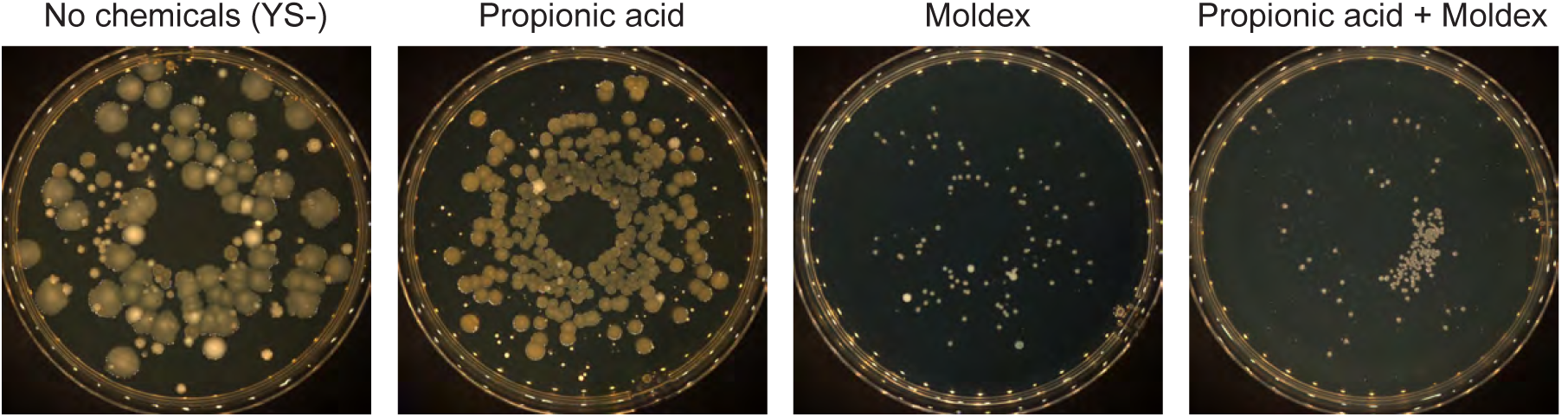
Alteration of the microbial diversity by preservatives contained in the fly diet. Wild-derived flies were kept on either a diet without chemical (YS-) or on the same diet supplemented with propionic acid, Moldex or both. After several generations, flies were crushed and the lysates plated on different culture media. Pictures show representative mannitol (Moldex diet and propionic acid + Moldex diet) or BHI (brain-heart infusion; YS- diet and propionic acid diet) plates and illustrate the decrease in microbial diversity caused by the introduction of chemicals in the fly diet, and more particularly the effect of Moldex.

### 3.3. Bacterial communities in the flies and in its diet are similar

To study the relationship between bacterial communities associated with flies and with their nutritive substrate, we used the wild-derived Drosophila population described above as a “natural” microbiota provider. The experiment was performed in duplicate, as described in the scheme on **Figure 3**. Statistical analyses showed that there was no significant effect of the experimental factors (the experimental repeat, the nature of the sample or the level in the protocol) on the alpha-diversity of the bacterial communities at both the order and the family level (**Table 2**). Only a weak three-way interaction of the factors was detected as statistically significant (p=0.047648) and only at the order level **(Table 2)**. We thus considered the effect of the experimental repeat on the bacterial composition as negligible and we decided to pool the results of the two technical replicates for further analyses as a means to increase the statistical power of our study.

**Figure 3.**
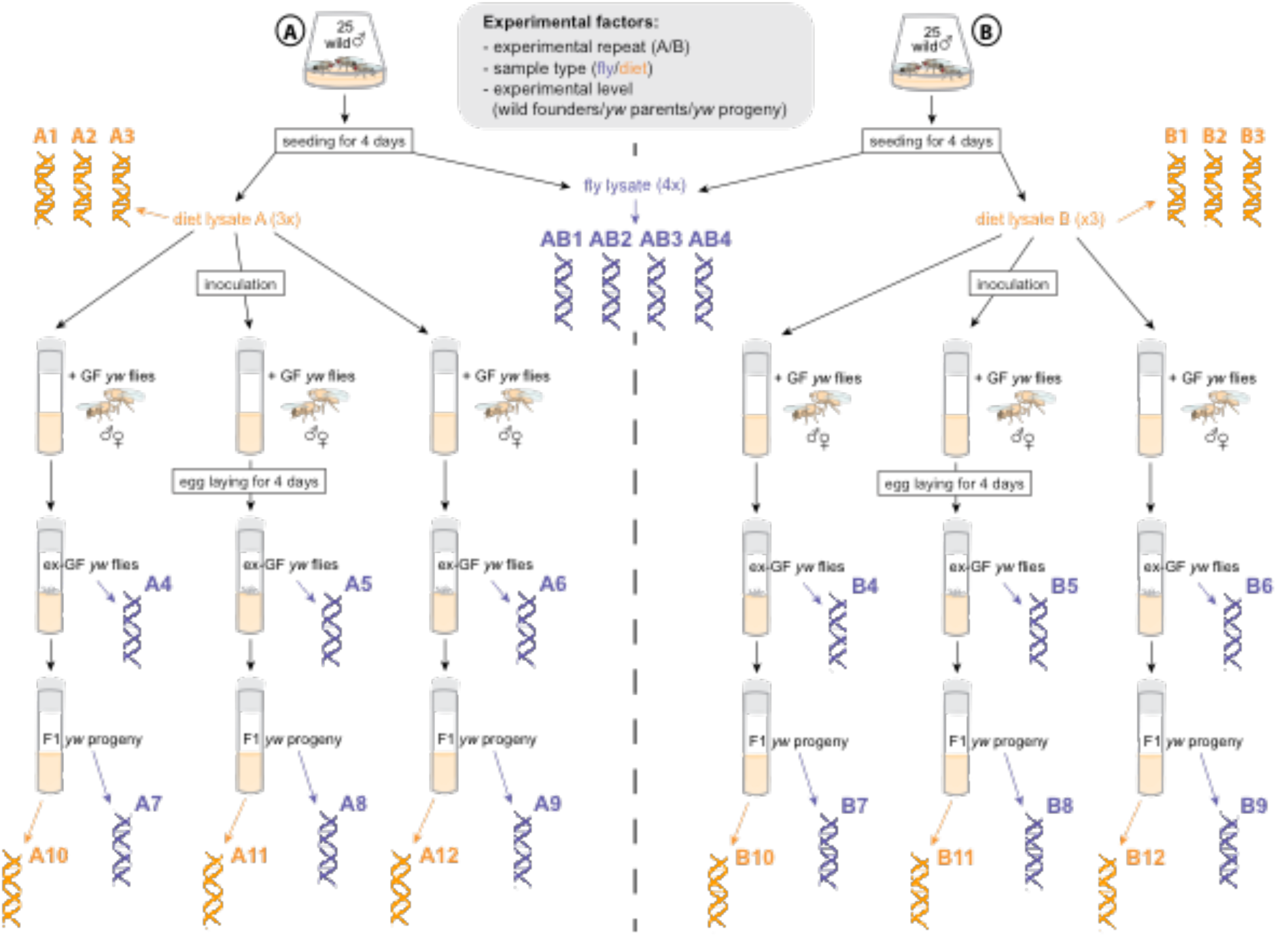
Schematic representation of the wild microbiota inoculation protocol and sample collection. Fly-derived DNA samples are represented in blue and diet-derived DNA samples are represented in yellow. The contribution of the experimental factors was statistically assessed with a type III ANOVA test. Such factors include the experimental repeat (replicate experiments A and B), the nature of the samples (fly samples and diet samples), as well as the experimental level (wild founders, *y,w* parents or *y,w* progeny). Note that the entire protocol was performed on YSexp diets.

**Table 2.** Type III ANOVA results for the Order and Family taxonomic level. Statistical analyses were performed on all samples from the wild-derived microbiota profiling experiment. Significant p-values (< 0.05) are shown in blue.

|  | Df | Sum of Sq | RSS | AIC | F value | Pr(>F) |
| --- | --- | --- | --- | --- | --- | --- |
| **Bacterial Order** |  |  |  |  |  |  |
| exp | 1 | 0 | 32 | 18 | 0 | 1 |
| sample.type | 1 | 0,190476 | 32,190476 | 18,189912 | 0,130952 | 0,720899 |
| level | 2 | 6,266667 | 38,266667 | 21,722985 | 2,154167 | 0,139837 |
| exp:sample.type | 1 | 1,523810 | 33,523810 | 19,488641 | 1,047619 | 0,317180 |
| exp:level | 2 | 4,916667 | 36,916667 | 20,57367096 | 1,690104 | 0,207588 |
| sample.type:level | 1 | 4,355556 | 36,355556 | 22,08355599 | 2,994444 | 0,097556 |
| exp:sample.type:level | 1 | 6,4 | 38,4 | 23,834290 | 4,4 | 0,047648 |
| **Bacterial Family** |  |  |  |  |  |  |
| exp | 1 | 7,82E-14 | 47,333333 | 30,527324 | 3,63E-14 | 1,000000 |
| sample.type | 1 | 0,047619 | 47,380952 | 30,559501 | 0,022133 | 0,883090 |
| level | 2 | 6,6 | 53,933333 | 32,704410 | 1,533803 | 0,237907 |
| exp:sample.type | 1 | 1,523810 | 48,857143 | 31,541270 | 0,708249 | 0,409081 |
| exp:level | 2 | 4,383333 | 51,716667 | 31,361414 | 1,018662 | 0,377487 |
| sample.type:level | 1 | 1,088889 | 48,422222 | 31,255134 | 0,506103 | 0,484306 |
| exp:sample.type:level | 1 | 3,6 | 50,933333 | 32,873014 | 1,673239 | 0,209242 |

Founder males were placed in cages for four days to seed the YSexp diet with the commensal bacteria they carried. As expected, after four days the bacteria had colonized this previously sterile diet. The presence of *Wolbachia* was detected only in the wild founder males and not on the diet **(Figure 4)**. This intracellular symbiont is indeed often found in fly populations, both in laboratory stocks and in the wild (Hoffmann et al. 1994; M. E. Clark 2005), and is transmitted vertically from mother to progeny. Its presence in the wild-derived population had been determined by PCR prior to this study **(Figure S2)**, and was confirmed with the 16S profiling data. Given its lifestyle and transmission mode, it is not surprising that *Wolbachia* is only found in the initial wild-derived fly samples and not in subsequent diet samples. Indeed, this protocol was based on a strict horizontal transmission of bacterial communities (i.e. via the diet). Consistently, the *y,w* stock was originally axenic and devoid of *Wolbachia,* therefore the endosymbiont was absent from the *y,w* parents and progeny samples. Since the experimental set-up induced a bias towards *Wolbachia* bacteria and prevented their transmission, the corresponding taxa (i.e. *Rickettsiales* order and *Anaplasmataceae* family) were excluded from all statistical analyses.

**Figure 4.**
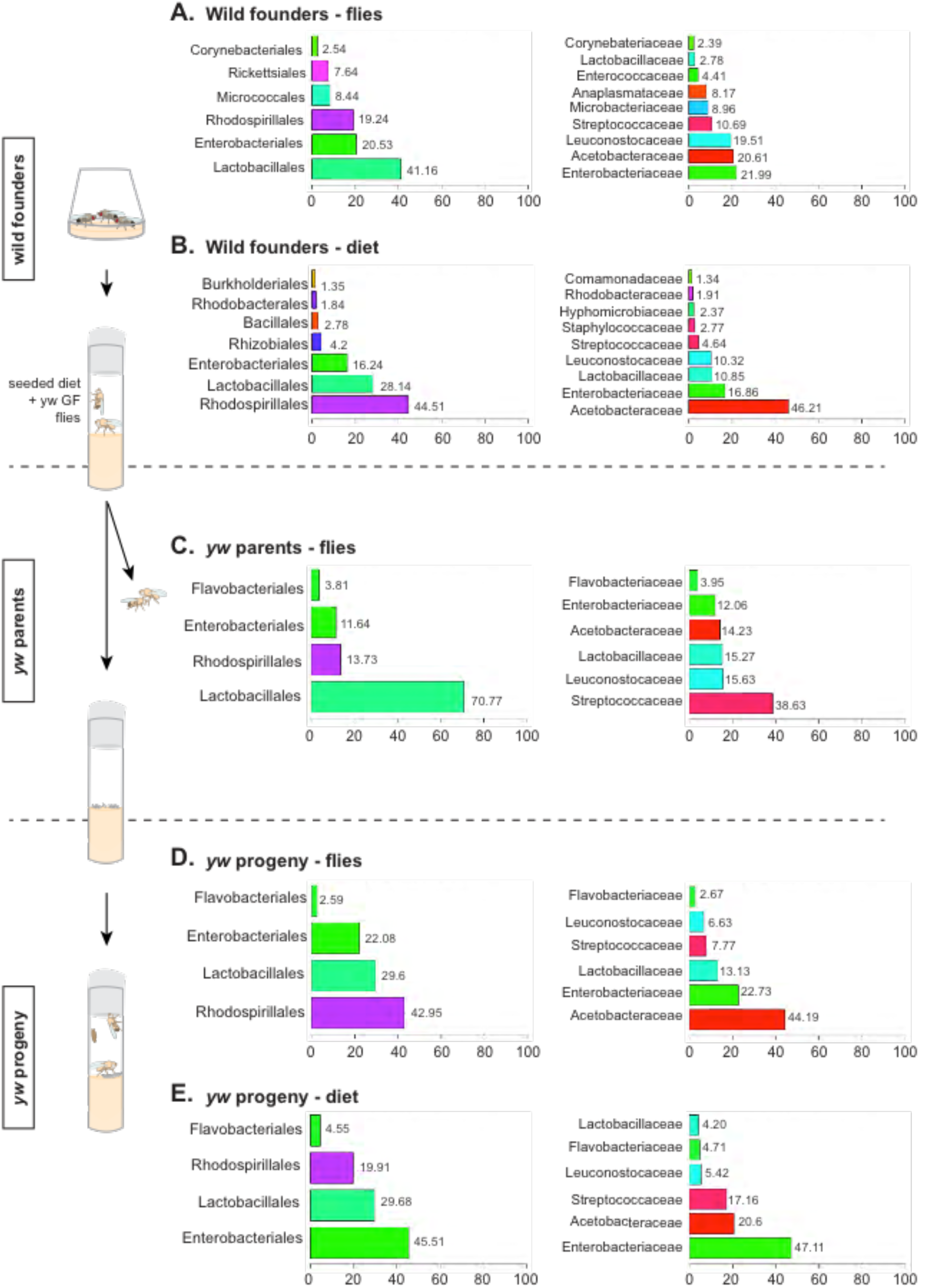
Bacterial communities associated with flies or diet upon transfer between populations and generations. Frequency of represented bacterial Orders and Families (left and right panels, respectively) in each sample group (pooled replicates) corresponding to the wild founder males (A, flies and B, diet), the *y,w* parents (C, flies) and the *y,w* progeny (D, flies and E, diet) levels. Equivalent percentages are indicated next to the corresponding bars. Graphs only represent the taxa present at a frequency of 1% or above.

In the founder male flies samples *Lactobacillales* were the most represented bacterial order, followed by *Enterobacteriales* and *Rhodospirillales* at similar frequencies **(Figue 4A, left panel)**. The hierarchy was different in the samples of YSexp diet seeded by these wild founders, with *Rhodospirillales* here being the main taxon **(Figure 4B, left panel)**. Additionally, within the *Lactobacillales* order several families were present and for some the ranking was modified between the wild founders flies and diet samples **(Figure 4A and B, right panels)**. Indeed, *Lactobacillaceae* that were originally ranked 8^th^ in the flies samples became the third most represented family in the diet samples. The position of the other members of the *Lactobacillales* order was either maintained (as for *Leuconostocaceae* and *Streptococcaceae)* or lowered (as for *Enterococcaceae* that fell below the cutoff of 1% of relative proportion in the diet samples).

The diet seeded by the wild-derived males was then used to inoculate fresh YSexp diet, and axenic (germ-free (GF); devoid of microbiota) *y,w* flies were placed on this bacteria-containing diet afterwards. After three days, the ex-GF flies were removed and their associated microbiota analyzed. Once again, the same three bacterial orders were found to dominate the community associated with these flies (*Lactobacillales, Rhodospirillales* and *Enterobacteriales* (**Figure 4C, left panel)**. Strikingly, even though directly originating from it, the hierarchy of the bacterial orders represented in the community associated with *y,w* parents was modified compared to the one of wild founders diet; here *Lactobacillales* were by far the most represented taxon. This was mainly due to a burst in the representation of bacteria from the *Streptococcaceae* family, which was the main taxon in the *y,w* parents samples. The second and third most represented families were the other *Lactobacillales* representatives from the wild founders diet samples, namely *Lactobacillaceae* and *Leuconostocaceae* **(Figure 4C, right panel)**.

After removal of their parents, the eggs laid by the ex-GF *y,w* flies were left to develop without manipulation, on the diet originally containing bacterial populations described in **Figure 4B**. When we analyzed the profile of the bacterial community associated with the *y,w* progeny, we found again the three same major orders, *Rhodospirillales, Lactobacillales* and *Enterobacteriales* **(Figure 4D, left panel)**. Here the ranking of the bacterial taxa was the same as that seen in the wild founders diet samples, with *Rhodospirillales* as the dominant order. In their associated diet however, the community was enriched in *Enterobacteriales* bacteria, which were now the main order represented **(Figure 4E, left panel)**. Since the *Rhodospirillales* and *Enterobacteriales* orders are represented each by a unique family in this study, it is unsurprising that the predominant bacterial families in the *y,w* progeny flies and diet samples were *Acetobacteraceae* and *Enterobacteraceae* **(Figure 4D and E, right panels)**.

Although the three major orders represented remained the same (namely *Enterobacteriales, Lactobacillales* and *Rhodospirillales*), the profile of the wild-derived bacterial community was slightly modified across samples through the experiment **(Figure 4)**. Among the represented taxa, the proportion of some bacterial orders was significantly different among the distinct sample types, notably the *Enterobateriales, Flavobacteriales* and *Lactobacillales* orders **(Figure 5A)**. This was further highlighted at the family level, with *Enterobacteraceae* (*Enterobacteriales*), *Flavobacteriaceae* (*Flavobacteriales*) and *Enterococcaceae* and *Streptococaceae* (*Lactobacillales*) being significantly differentially represented through the experiment **(Figure 5B)**. However these observations do not correspond to a pattern that would support the specific association of particular bacterial taxa with a given type of sample. Furthermore, as previously mentioned, the global statistical analyses performed on all experimental samples showed no significant effect of either the nature (fly/diet) or the level (wild founders/*y,w* parents/*y,w* progeny) of the samples **(Table 2)**. This was further confirmed when we carried out principal component analyses to cluster samples according to the similarities in their community profile both at the order and family levels (**Figure 6A-C left and right panels** respectively). Indeed, such analyses revealed that neither the type of sample, the level of the experiment or the experimental replicate considered could explain the differences observed among the samples **(Figure 6 and Table 3)**.

**Figure 5.**
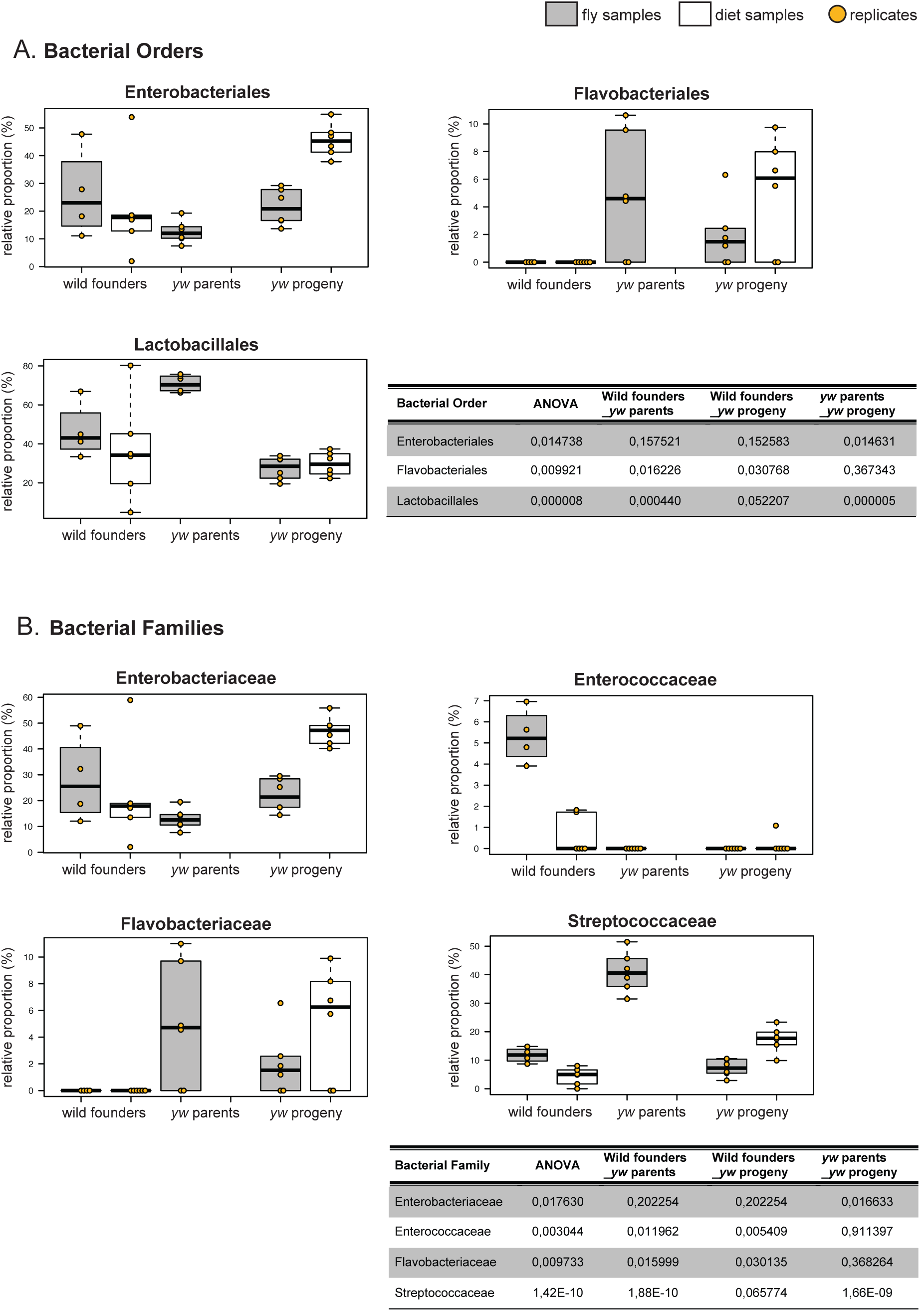
Significantly differentially represented bacterial taxa. The proportion of each bacterial Order (A) and Family (B) with significantly different representation across samples is given for each combination of sample type (fly/diet) and experimental level (wild founders/*y,w* parents/*y,w* progeny). The results of the statistical analyses are summarized in the corresponding tables, with p-values of the type III ANOVA and subsequent pairwise-t-tests between modalities.

**Figure 6.**
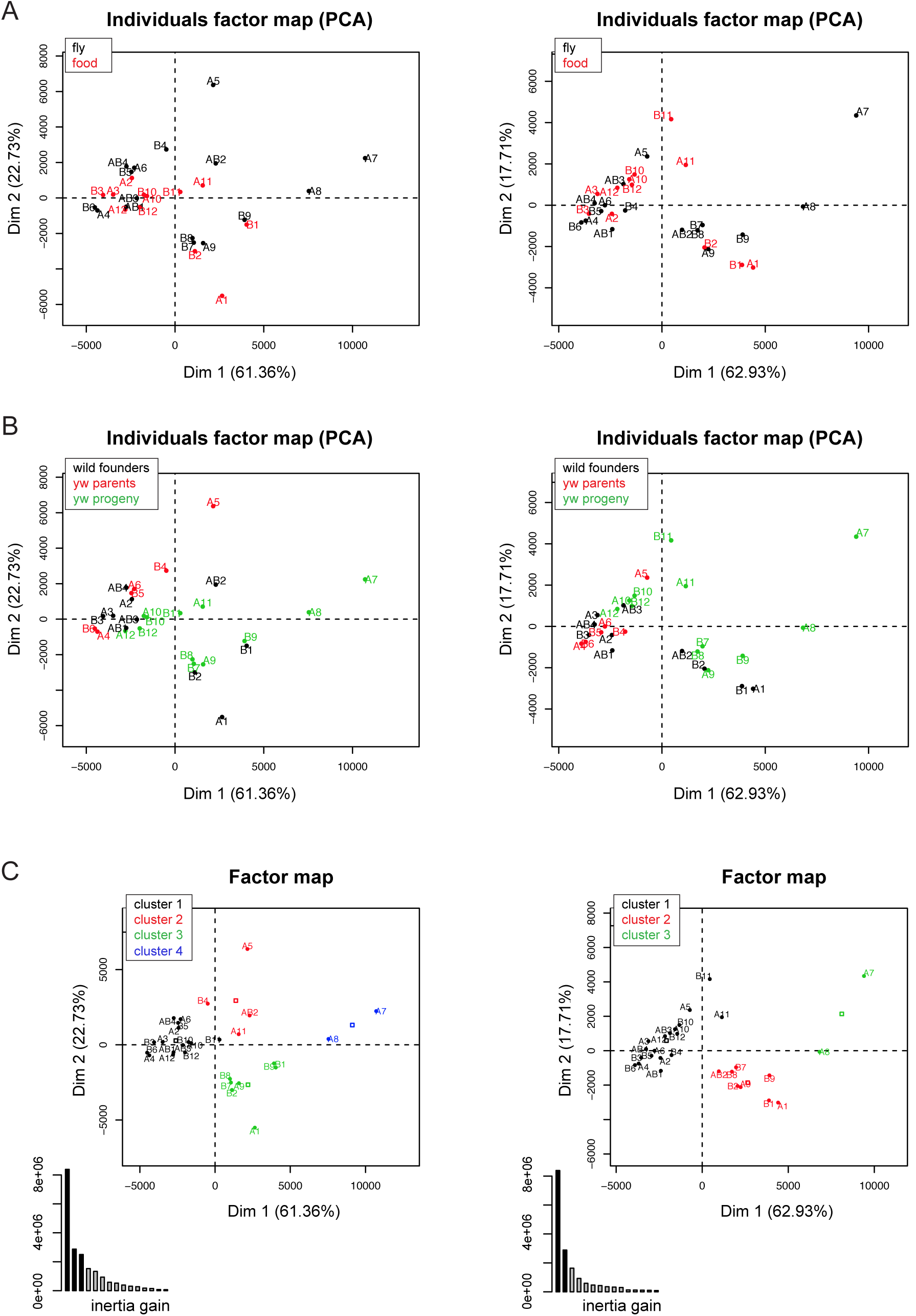
Differences in composition among bacterial communities are not explained by the type of sample or the experimental level. Principal component analyses results are shown for the bacterial Orders and Families (left and right panels, respectively). Samples have been color-coded according to their type (A, fly/diet) or their experimental level (B, wild founders/*y,w* parents/*y,w* progeny). Panels C show in different colors the clusters resulting from the hierarchical clustering on principal components, together with the corresponding inertia gain graphs.

**Table 3.** Chi2 test results for the Order and Family taxonomic level. Statistics were performed on the results of the hierarchical clustering on principal components analyses.

|  | p.value | df |
| --- | --- | --- |
| **Bacterial Order** |  |  |
| level | 0,308393167 | 6 |
| experiment | 0,379903741 | 6 |
| sample.type | 0,44140872 | 3 |
| **Bacterial Family** |  |  |
| level | 0,162702193 | 4 |
| sample.type | 0,373672699 | 2 |
| experiment | 0,386327304 | 4 |

## 4. Discussion & Conclusions

In order to determine if our reference *yellow white* stock could serve as a microbiota-provider, we first analyzed the composition of the bacterial community associated with this fly population and found it to be poorly diverse. Besides *Wolbachia,* the *y,w* stock was associated with only three bacterial genera (*Corynebacterium, Enterobacter* and *Lactobacillus*), and among them, the *Lactobacillus* genus was by far the most represented. This result was reminiscent of those published by Sharon and colleagues who reported the effect of the microbiota on assortative mating. In this particular study, switching flies from a molasses-based diet to a starch-based diet dramatically impacted the bacterial diversity associated with their fly stock, which ended up becoming mono-associated with *Lactobacillus plantarum* (Sharon et al. 2010). In our case, the *y,w* stock tested is also reared on a starch-based diet, since cornmeal is the main carbohydrate source in our standard fly diet, which might at least in part explain the low bacterial diversity. Moreover, bacteria from the genus *Acetobacter* (another taxa known as a major fly commensal) thrive mainly on simple sugars, which could explain their absence from this community. Another feature of the fly diet composition could also explain the reduced bacterial diversity of the microbiota associated with our *y,w* stock: in addition to being starch-based, the diet routinely used in our laboratory contains Moldex and propionic acid, two chemicals commonly added as preservatives in fly diet recipes. The very purpose of these chemicals is to prevent diet spoilage by antagonizing microbial development (mostly fungal). In this light it is anticipated that they might also hinder the growth of commensal bacteria, thus reducing the microbial diversity of laboratory-reared fly stocks.

To perform this bacterial dynamics study, we aimed to start with as much bacterial diversity as possible. First because we thought it would increase the chances to observe potential shifts in the community composition, and secondly to make our setup closer to a natural setting. Indeed, it has been previously shown that the diversity of wild populations’ microbiota was increased compared to that of laboratory-reared flies (Chandler et al. 2011; Staubach et al. 2013). We thus decided to turn to the wild to find our starting microbial community. This population allowed us to confirm the adverse effect of Moldex and propionic acid on commensal bacterial diversity. Furthermore, in our setup, Moldex seemed to have the strongest impact on microbiota composition and drastically reduced the diversity of microbial communities associated with the fly population. We therefore designed a diet deprived of Moldex and containing a reduced amount of propionic acid for our study in order to sustain diversity of the fly microbiota.

As shown by the 16S-based profiling of their associated community, when kept in the laboratory on our diet designed to favor bacterial diversity (YS- diet), our wild-derived fly population indeed harbored a much more complex community than that of the *y,w* stock; even if the proportions were not strictly maintained, almost all bacterial taxa present in the wild founder males were transferred to the diet and to the *y,w* flies. In the graphical representation of the community profiles, we arbitrarily chose to represent only the taxa with a frequency proportion of 1% or above. This cutoff explains why certain taxa appearing in the wild founder males graphs are not present anymore in those describing the *y,w* samples; their proportion dropped below the 1% cutoff, hence their absence from the graphs **(Figure 4)**. One bacterial type, the *Flavobateriales* order, was however enriched in the *y,w* samples. Indeed, in the wild founder flies and diet samples *Flavobateriales* were underrepresented, with proportions of around 0.3% and 0.2% respectively while this proportion increased significantly in all *y,w* samples; *Flavobacteriales* were systematically found among the 4 most represented taxa, and above the 1% cutoff. Nevertheless, the 3 main orders always represented were the *Enterobacteriales, Lactobacillales* and *Rhodospirillales,* which were previously reported as major components of the fly microbiota. Indeed, even if the representative genera tend to vary across studies and populations, these higher taxonomic levels are almost always present (Chandler et al. 2011; Staubach et al. 2013).

There was no significant distinction in the composition of the communities between the fly and the diet samples; no bacterial taxa were more, or singularly represented in one type of samples. Thus, the flies did not appear to actively select for or against certain bacteria, and the bacterial content of the flies was similar to that of the nutritive substrate. Nonetheless, this point could only be fully assessed by performing a complementary experiment, with the same initial set up of diet seeded by the wild founder flies, but without adding any flies subsequently. Such an experiment would allow us to study the dynamics of the bacterial community in a fly-free environment and to confirm the absence of effect of the flies on their microbiota composition. Interestingly in a recent study, Wong and colleagues performed this type of experiment and found that the presence of Drosophila promoted the maintenance of *Lactobacillus* bacteria in the niche. Furthermore, the presence of the fly partially protected *Lactobacilli* against the antagonistic effect of *Acetobacter,* which in a fly-free environment are taking over and in turn completely dominate the community (Wong et al. 2015). Surprisingly however, these promoting and protective effects were observed only when flies were present at high densities, suggesting that whatever the underlying cause of this impact, the fly population needs to attain a critical mass to exert it. Additionally, such effects were revealed in a set of experiments using a poorly diverse microbiota, consisting of an artificial mixture of four bacterial species all belonging to the *Acetobacter* and *Lactobacillus* genera. These bacterial strains were originally derived from laboratory-reared fly stocks (Wong et al. 2013) and might have adapted to laboratory conditions and to manufactured fly diets. Their behavior might thus be very different from the dynamics of a natural microbiota.

For the majority of the bacterial orders identified in this study, only one family was represented for each order. The *Lactobacillales* were the only exception, as *Lactobacillaceae, Leuconostocaceae* and *Streptococcaceae* families were represented **(Figure 4)**. Unsurprisingly, the relative proportions of the represented families followed those of their respective orders. We also analyzed the results at the genus level and drew the same conclusion (data not shown). The genera results are however less representative in this study since a considerable proportion of sequences were unclassified at this taxonomic level and thus excluded from our analysis. We cannot rule out that potential shifts in the community structure and composition could only be revealed at this taxonomic level or at the species one. Furthermore, it has been shown that different strains of the same bacterial species could have very different impacts (Storelli et al. 2011; Chaston et al. 2014), which highlights the importance of considering this taxonomic level as well. However, our 16S-based community profiling protocol is not amenable to such in-depth discrimination of bacterial taxa. Additionally, our study is restricted to the analysis of the first-degree progeny. It might be of interest to conduct such an analysis for a longer period of time and observe how the bacterial community dynamics changes with additional generations.

All together, our results indicate that the microbiota of Drosophila is stably associated with the fly population and its nutritional substrate within a given environmental niche, and upon transfer between populations and generations. Adult flies seed their nutritive substrate with the microorganisms they initially carry, allowing the microbial community to establish and invade the niche. This microbiota thus remains associated with the fly population occupying this shared habitat and is transmitted to their progeny. At each generation, freshly hatched larvae associate and develop with the microbiota and each individual will later on facilitate the dispersal of their associated bacteria. Adult Drosophila will then migrate to new habitats, bringing along all or part of the initial bacterial community and seed new niches. This study thus illustrates the stable association of a Drosophila-derived microbiota with both its animal partner and the nutritional environment and indicate that the nutritional substrate is an important microbiota habitat to integrate in Drosophila/microbiota studies.

## 5. Annexes

### 5.1. Acknowledgments

The authors would like to thank the Arthro-Tools platform of the SFR Biosciences (UMS3444/US8) for providing Drosophila husbandry materials, the IGFL sequencing platform for performing the deep sequencing experiments, Loan Bozonnet for fly diet preparation and Dali Ma for critical reading and edition of the manuscript.

### 5.3. Authors contribution

FL supervised the work. MT and FL designed the experiments and MT performed them. BG and SH designed the 16S-based profiling protocols. BG prepared the sequencing libraries and performed the sequencing. PJ and SH assisted MT for BioIT and BioStats analyses. MT, PJ, SH and FL analysed the results. MT wrote the manuscript with inputs from FL.

### 5.4. Funding

This work was funded by an ERC starting grant (FP7/2007-2013-N°309704). The lab is supported by the FINOVI foundation, the “Fondation Schlumberger pour l'Education et la Recherche” and the EMBO Young Investigator Program.

---

**Figure S1.**
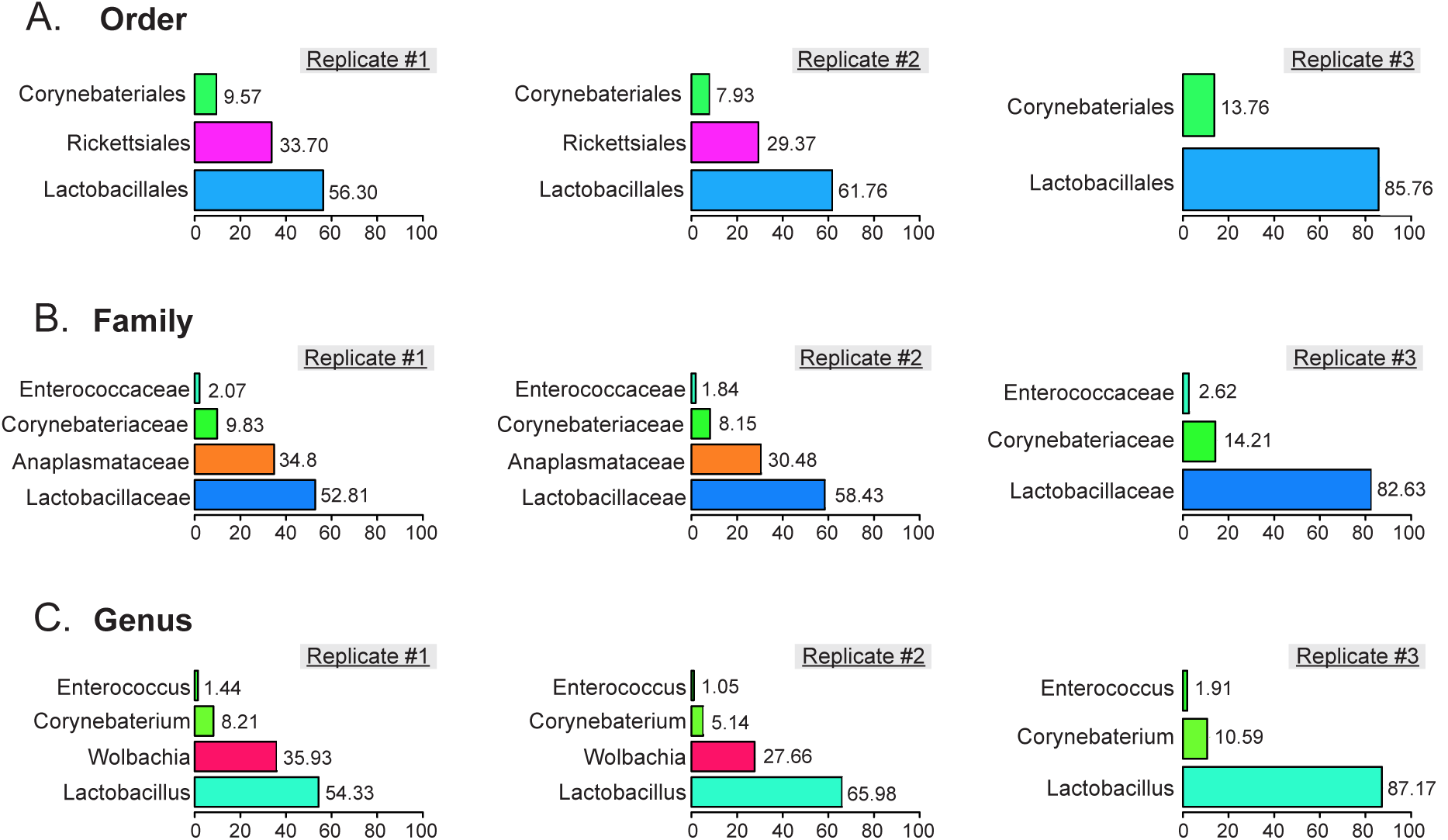
Community profile of the *yellow white* stock microbiota. Frequency of bacterial taxa present in each of the three biological replicates. Taxonomic levels used: Order (A), Family (B) and Genus (C). Equivalent percentages are indicated next to the corresponding bars. Graphs only represent the taxa present at a frequency of 1% or above.

**Figure S2.**
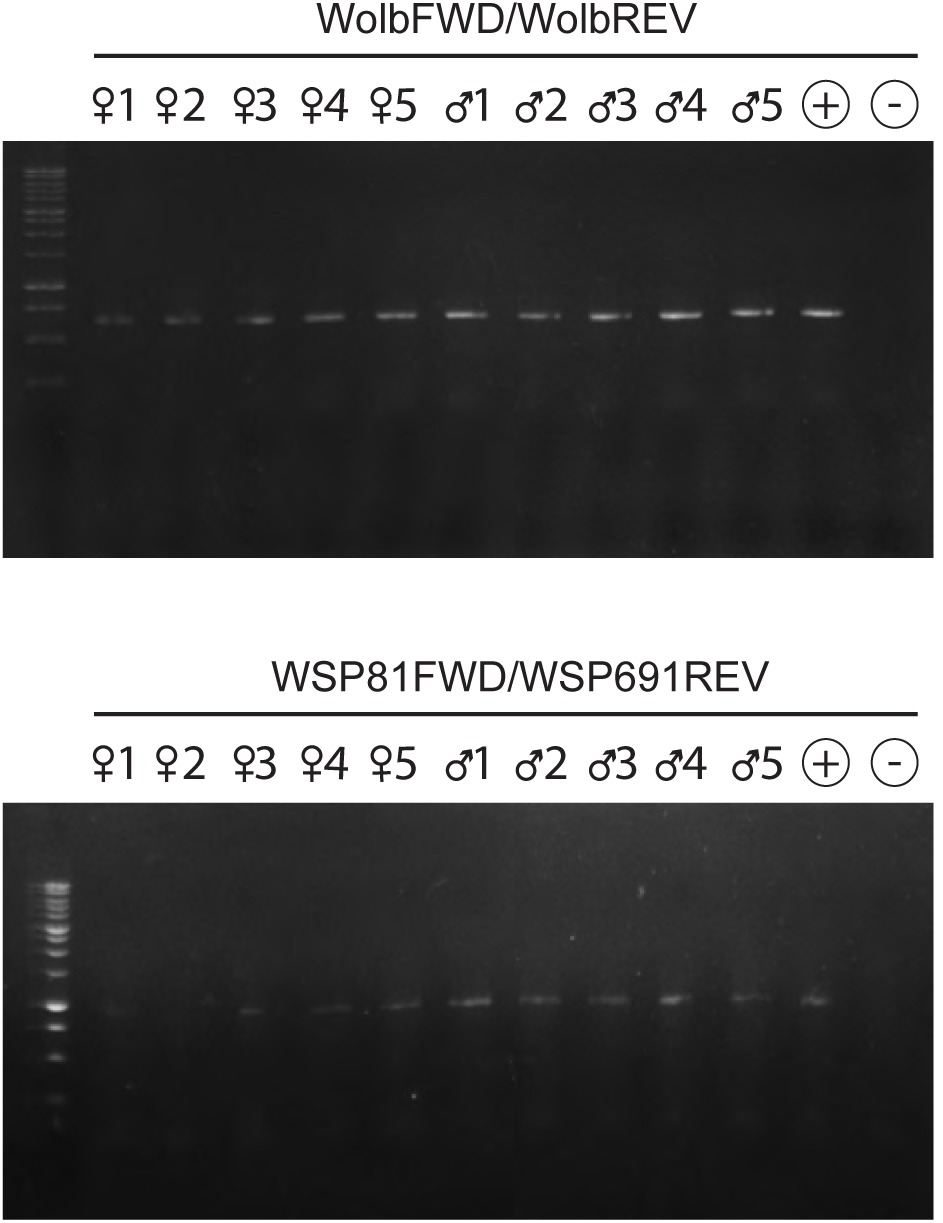
The wild-derived Drosophila population is infected with Wolbachia. Positive PCR result for Wolbachia presence. The PCR amplification was performed with general (upper gel) and strain-specific (lower gel) primer pairs.

## References

Broderick, N.A. & Lemaitre, B., 2012. Gut-associated microbes of *Drosophila melanogaster*. Gut Microbes, 3(4), pp.307–321.

Brummel, T. et al., 2004. Drosophila lifespan enhancement by exogenous bacteria. Proceedings of the National Academy of Sciences of the United States of America, 101(35), pp.1297–12979.

Chandler, J.A. et al., 2011. Bacterial communities of diverse Drosophila species: ecological context of a host-microbe model system. PLoS Genetics, 7(9), p.e1002272.

Chaston, J.M., Newell, P.D. & Douglas, A.E., 2014. Metagenome-wide association of microbial determinants of host phenotype in *Drosophila melanogaster*. mBio, 5(5), pp.e01631–14.

Clark, M.E., 2005. Widespread Prevalence of Wolbachia in Laboratory Stocks and the Implications for Drosophila Research. Genetics, 170(4), pp.1667–1675.

Clark, R.I. et al., 2015. Distinct Shifts in Microbiota Composition during Drosophila Aging Impair Intestinal Function and Drive Mortality. Cell Reports, 12(10), pp.1656–1667.

Erkosar, B. et al., 2013. Host-Intestinal Microbiota Mutualism: "Learning on the Fly''. Cell Host & Microbe, 13(1), pp.8–14.

Guo, L. et al., 2014. PGRP-SC2 Promotes Gut Immune Homeostasis to Limit Commensal Dysbiosis and Extend Lifespan. Cell, 156(1-2), pp.109–122.

Hamady, M. et al., 2008. Error-correcting barcoded primers for pyrosequencing hundreds of samples in multiplex. Nature Methods, 5(3), pp.235–237.

Hoffmann, A.A., Clancy, D.J. & Merton, E., 1994. Cytoplasmic incompatibility in Australian populations of Drosophila melanogaster. Genetics, 136(3), pp.993–999.

Kohler, R.E., 1994. Lords of the Fly: Drosophila Genetics and the Experimental Life, The University of Chicago Press.

Ma, D. et al., 2015. Studying host-microbiota mutualism in Drosophila: Harnessing the power of gnotobiotic flies. Biomedical Journal, 38(4), pp.285–293.

Martin, M., 2011. Cutadapt removes adapter sequences from high-throughput sequencing reads. EMBnet.journal, 17(1), p.10.

McFall-Ngai, M. et al., 2013. Animals in a bacterial world, a new imperative for the life sciences. In Proceedings of the National Academy of Sciences of the United States of America. pp. 3229–3236.

Petkau, K. et al., 2014. A Deregulated Intestinal Cell Cycle Program Disrupts Tissue Homeostasis without Affecting Longevity in Drosophila. Journal of Biological Chemistry, 289(41), pp.28719–28729.

Ryu, J.H. et al., 2008. Innate Immune Homeostasis by the Homeobox Gene Caudal and Commensal-Gut Mutualism in Drosophila. Science, 319(5864), pp.777–782.

Salter, S.J. et al., 2014. Reagent and laboratory contamination can critically impact sequence-based microbiome analyses. BMC Biology, 12(1), p.118.

Sang, J.H., 2001. Drosophila melanogaster: the fruit Fly E. C. R. Reeve, ed. Encyclopedia of Genetics.

Schloss, P.D. et al., 2009. Introducing mothur: open-source, platform-independent, community-supported software for describing and comparing microbial communities. Applied and Environmental Microbiology, 75(23), pp.7537–7541.

Sharon, G. et al., 2010. Commensal bacteria play a role in mating preference of Drosophila melanogaster. Proceedings of the National Academy of Sciences of the United States of America, 107(46), pp.20051–20056.

Shin, S.C. et al., 2011. Drosophila Microbiome Modulates Host Developmental and Metabolic Homeostasis via Insulin Signaling. Science, 334(6056), pp.670–674.

Staubach, F. et al., 2013. Host Species and Environmental Effects on Bacterial Communities Associated with Drosophila in the Laboratory and in the Natural Environment. PLoS ONE, 8(8), p.e70749.

Storelli, G. et al., 2011. *Lactobacillus plantarum* Promotes Drosophila Systemic Growth by Modulating Hormonal Signals through TOR-Dependent Nutrient Sensing. Cell Metabolism, 14(3), pp.403–414.

Téfit, M.A. & Leulier, F., 2017. *Lactobacillus plantarum* favors the early emergence of fit and fertile adult Drosophila upon chronic undernutrition. Journal of Experimental Biology.

Venu, I. et al., 2014. Social attraction mediated by fruit flies' microbiome. Journal of Experimental Biology, 217(8), pp.1346–1352.

Wang, Y. & Qian, P.-Y., 2009. Conservative Fragments in Bacterial 16S rRNA Genes and Primer Design for 16S Ribosomal DNA Amplicons in Metagenomic Studies D. Field, ed. PLoS ONE, 4(10), p.e7401.

Wong, A.C.-N. et al., 2015. The Host as the Driver of the Microbiota in the Gut and External Environment of Drosophila melanogaster. Applied and Environmental Microbiology, 81(18), pp.6232–6240.

Wong, A.C.-N., Chaston, J.M. & Douglas, A.E., 2013. The inconstant gut microbiota of Drosophila species revealed by 16S rRNA gene analysis. The ISME Journal, pp.1–11.

Wong, A.C.N., Dobson, A.J. & Douglas, A.E., 2014. Gut microbiota dictates the metabolic response of Drosophila to diet. Journal of Experimental Biology, 217(11), pp.1894–1901.

